# New insights on the role of *SlDMR6-1* in drought avoidance in tomato

**DOI:** 10.1101/2023.12.14.571645

**Authors:** Maioli Alex, De Marchi Federica, Valentino Danila, Gianoglio Silvia, Patono Davide, Miloro Fabio, Bai Yuling, Comino Cinzia, Lanteri Sergio, Lovisolo Claudio, Acquadro Alberto, Moglia Andrea

**Author notes:** Corresponding author: Moglia Andrea.

## Abstract

The DOWNY MILDEW RESISTANCE 6 (DMR6) protein is a 2-oxoglutarate (2OG) and Fe(II)-dependent oxygenase, involved in salicylic acid (SA) metabolism, and its inactivation in tomato was found to increase SA levels and to confer disease-resistance against several pathogens. SA is also recognized as an abiotic stress-tolerance enhancer, and we tested the resistance to drought stress in *Sldmr6-1* tomato mutants generated by the CRISPR/Cas9 technique.

Wild-type (WT) tomato cultivar ‘San Marzano’ and its *Sldmr6-1* mutants were subjected to water deprivation for 7 days. At the end of the period, while WT plants exhibited severe wilting, the T_2_ *Sldmr6-1* mutant plants showed turgid leaves and maintained higher Soil Relative Water Content (SRWC). *Sldmr6-1* mutants adopted a water saving behaviour reducing transpiration rate (E) by decreasing stomatal conductance (Gs). Assimilation rate (A) decreased in parallel to E under drought stress, resulting in no alteration of the CO_2_ concentration in the sub-stomatal chamber (Ci) and increasing the Water Use Efficiency (WUE, A/E). Defence mechanisms of the photosynthetic machinery triggered in *Sldmr6-1* mutants, that under drought stress showed up-regulation of the genes *SlAPX* and *SlGST* (anti-oxidant related) as well as down-regulation of *SlCYP707A2* gene, which is involved in ABA catabolism. Our results suggest that the disabling of *SlDMR6-1* in tomato plants leads to a drought-avoidance strategy through tight control of stomatal closure controlling water loss. In addition, it was highlighted, for the first time in tomato, that *Sldmr6-1* mutants showed reduced susceptibility to *Phytophthora infestans*, the causal agent of Late Blight.

## INTRODUCTION

Rising temperatures and shifts in precipitation patterns pose a serious concern in various parts of the world, leading to a substantial reduction in the availability of water during the crop cultivation season. Drought stress is a limiting factor of plant growth and concurs with plant pathogen or pest attacks to cause severe plant yield reductions (Cappetta et, 2020). World food security in the coming years will hence largely depend on the availability of biotic and abiotic stress-tolerant plants. In the past, research has been focused on elucidating the mechanisms involved in the response to individual stress, and more recently on identifying new forms of resistance to multiple stress, and several genes participating in recognizing both biotic and abiotic stresses have been characterized (Saijo et al, 2020; Sunarti et, 2022). Tomato (*Solanum lycopersicum* L.) suffer severe yield losses due to both abiotic and biotic stresses (Kissoudis et al, 2016), thus the development of elite genotypes endowed of tolerance towards them is a major objective for tomato breeders (Egea et al, 2022).

To this end, a significant contribution can be provided by the emergent CRISPR/Cas9 technology for genome editing, which may greatly contribute to precision breeding and reducing product development costs compared to complex, imprecise and lengthy conventional breeding strategies (Lassoued et al, 2019; Zhu et al, 2020). Genome editing has been applied to tomato since 2014 (Brooks et al, 2014; Lor et al, 2014) and has greatly facilitated the functional characterization of genes involved in fruit yield and quality, development and ripening, stress response and domestication processes (Vu et al, 2020; Lobato-Gómez et al, 2021; Salava et al, 2021).

Negative regulators of abiotic stress response pathways have been targeted in tomato through CRISPR/Cas9, and key genes involved in drought (*SlLBD40, SlMAPK3, SlNPR1, SlARF4, SlGID1a*), salinity (*SlARF4, SlHyPRP1*) and chilling stress (*SlCBF1, SlBZR1*) response have been identified (Salava et al, 2021; Bouzroud et al, 2020; Tran et al, 2021; Yin et al, 2018).

One mechanism of pathogen plant resistance is due to the loss-of-function of genes required for the onset of pathogenesis, referred to as plant susceptibility (S) genes (Pavan et al, 2010), which have been proposed as key targets for genome editing approaches (Engelhardt et al, 2018; Chaudary et al, 2022). A classic example of a S-gene is Mildew resistance locus O (*Mlo1*) (Bü et al, 1997), of which loss-of-function natural mutants have been exploited for over 70 years in barley breeding programs (Piffanelli et al, 2002). In tomato, successful examples of CRISPR/Cas9 genome editing aimed at disabling S-genes have been reported to confer resistance against distinct classes of pathogens like viruses (Atarashi et al, 2020; Yoon et al, 2020; Kuroiwa et al, 2022), bacteria (Ortigosa et al, 2019; De Toledo Thomazella et al, 2021), fungi (Nekrasov er al, 2017; Santillán Martínez et al, 2020) and oomycetes (De Toledo Thomazella et al, 2021; Li et al, 2022).

Among S-genes, DOWNY MILDEW RESISTANCE 6 (*DMR6*) is particularly interesting. It encodes a 2-oxoglutarate (2OG) and Fe(II)-dependent oxygenase, which reduces the active salicylic acid (SA) pool acting as SA 5-hydroxylase (Van Damme et al, 2008; Zhang et al, 2017). CRISPR/Cas9 knock-out mutants of *DMR6* have been generated in different species such as *Arabidopsis thaliana* (Zeilmaker et al 2015,), *Vitis vinifera* (Giacomelli et al, 2022), *Ocimum basilicum* (Hasley et al, 2021), *Musa spp*. (Tripathi et al, 2021), *Solanum tuberosum* (Kieu et al, 2021), and *Citrus spp*. (Parajuli et al, 2022).

In tomato, impairment of *SlDMR6-1* has resulted in resistance to bacteria (*Pseudomonas syringae, Xanthomonas gardneri, Xanthomonas perforans)*, oomycetes (*Phytophthora capsici)* and fungi (*Pseudoidium neolycopersici)* (De Toledo Thomazella et al, 2021). According to Kieu et al (2021), the disabling of the potato *DMR6-1* gene resulted in plants with increased resistance to Late Blight (LB) caused by *Phytophthora infestans*. Remarkably, enhanced SA levels and transcriptional activation of immune response correlates with disease resistance (De Toledo Thomazella et al, 2021).

SA is a natural phenolic compound and a signalling regulator, which plays various regulatory roles in mediating plant development, growth, and defence to environmental stresses. SA has been shown to improve plant tolerance to major abiotic stresses such as salinity, metal, osmotic, drought and heat stresses (Khan et al, 2015; Liu et al, 2022). Furthermore, it has been observed that exogenous application of SA in low concentration in tomato could mitigate the oxidative stress generated by the water stress (Chakma et al, 2021; Aires et al, 2022). Interestingly, tomato plants pre-treated with SA and experiencing water deficit had an improved water-use efficiency and net photosynthetic rate, leading to an increased fruit weight, fruit numbers, and biomass (Lobato-Gómez et al, 2021).

In this study, CRISPR/Cas9 editing technology was used for disabling the *SlDMR6-1* gene in the tomato ‘San Marzano’ variety, an Italian local landrace used in the canning industry. Through Illumina Whole Genome Sequencing (WGS) we assessed potential off-target effects and mutational status of one selected T_1_ mutant, characterized by *Cas9* absence. Due to the link between SA and drought stress response, we tested the potential drought resistance of *Sldmr6-1* tomato mutants. In addition, we characterized their tolerance to Late blight, a potentially devastating disease of tomato.

## RESULTS

### Molecular screening of *Sldmr6-1* mutants

A CRISPR-Cas9 vector containing the h*Cas9* gene, the selective marker (*NptII*) and the polycistronic tRNA–gRNA structure with the 3 gRNAs targeting the *SlDMR6-1* gene was introduced via *Agrobacterium tumefaciens*-mediated transformation into the tomato cultivar ‘San Marzano’. These gRNAs target the first three exons of *SlDMR6-1* in order to disrupt the protein’s catalytic site (Fig.S1). Editing efficiency spanned greatly between the targets in T_0_ plants: it was higher for gRNA1 and gRNA3, while no editing was detectable for gRNA2 (Data S1).

T_1_ plants were sequenced by Sanger approach at the target loci to evaluate editing efficiency and transmission pattern of CRISPR/Cas9-induced mutations using the web tool TIDE (Brinkman et al, 2018). Out of the 14 analysed individuals, 6 showed homozygous, heterozygous and biallelic mutations at the target sites of gRNA1 and gRNA3 (Fig.1; Data S1). The T_1__6 plant showed homozygous mutations for gRNA1 (-3/-3) and gRNA3 (+1/+1), and the presence of h*Cas9*. The T_1__7 plant was then selected for further molecular analyses due to both its homozygous mutations for gRNA1 (+1/+1) and gRNA3 (+1/+1), and the absence of any transgene.

**Fig. 1.**
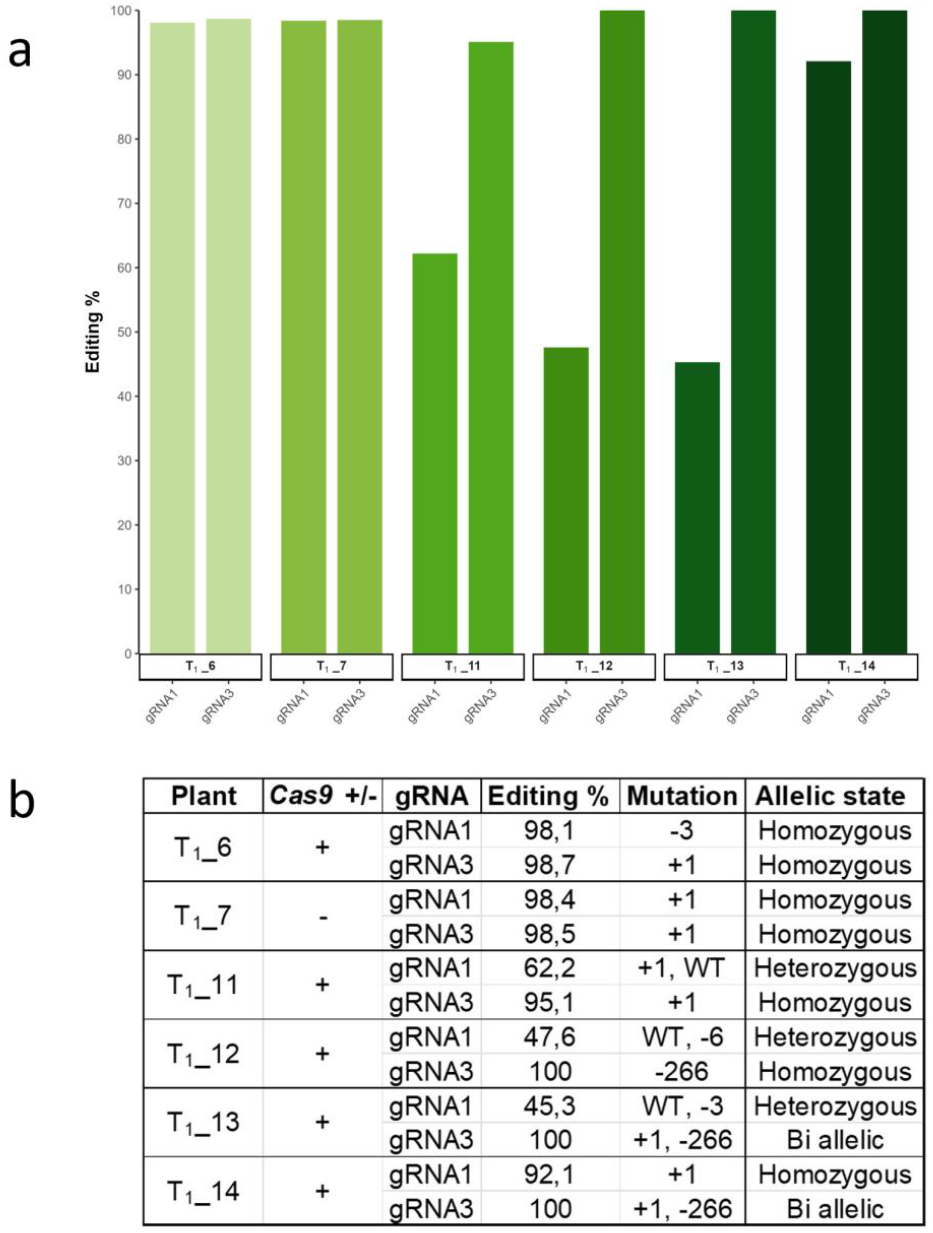
Genotyping of targeted gene mutations induced by CRISPR/Cas9 in selected T_1_ plants. (a) Mutagenesis frequencies (%) at gRNA1 and gRNA3 targets in six plants of the T_1_ progenies. (b) h*Cas9* presence, observed mutations and allelic forms. Data were retrieved through TIDE analysis of Sanger sequences.

### Whole genome resequencing of a *Sldmr6-1* mutant

T_1__7 and WT plants were sequenced through Illumina WGS. Genome sequencing of T_1__7 yielded 196,4 million raw paired-end reads (29.5 Gb), with an average length of 150 bp. These were reduced to 196,1 million after filtering and trimming high-quality reads. The sequence depth of coverage ranged from 37.7X (T_1__7) to 42.8X (WT) (Data S1).

A *de novo* genome assembly of T_1__7 was produced and the integration of T-DNA was inspected through the scanning of the scaffolds with Blast analysis, which did not identify any T-DNA insertions. These results clearly demonstrated h*Cas9* segregation.

Scanning of *SlDMR6-1* in the gRNA1 region revealed a 100% editing effect with homozygous mutations (a 1 bp insertion at position SL4.0ch03:46628534) and no reference alleles, supporting the analysis performed with TIDE (Fig.2). Scanning of *SlDMR6-1* in the gRNA3 region highlighted an editing efficiency of 100% with homozygous mutations (a 1 bp insertion at position SL4.0ch03:46624776) and no reference alleles, in agreement with the TIDE analysis (Fig.1a and b). The mutations impacting *SlDMR6-1* result in a premature stop codon in exon 1 leading to a truncated protein.

**Fig. 2.**
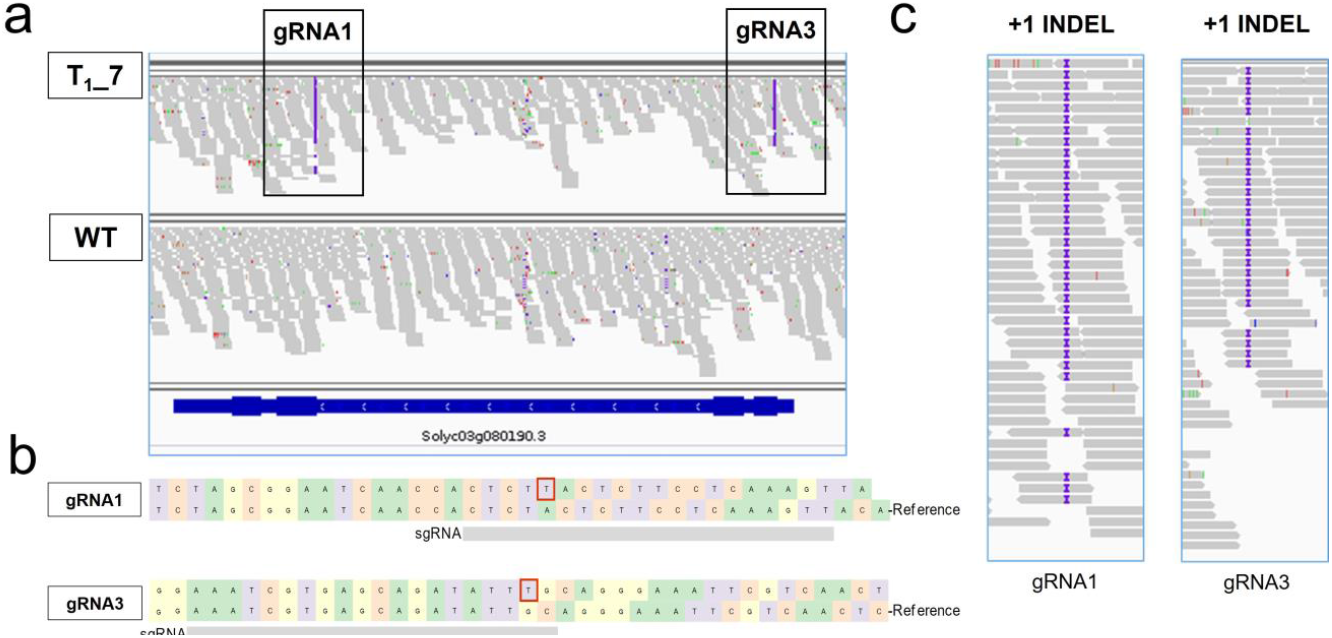
Comprehensive Whole Genome Sequencing of T1_7 *Sldmr6-1* mutant. (a) Detailed sequence alignment depicting the edited *SlDMR6-1* gene and its alignment to the two sgRNAs (gRNA1 and gRNA3) within both the T1_7 plant and the WT plant. (b) Genotyping analysis illustrating the targeted gene mutations in the T1_7 plant obtained with CRISPR/Cas9. (c) Focus on sgRNA1 and sgRNA3, displayed on the right-hand side.

### Off-target and SNP analyses in *Sldmr6-1* mutant

To confirm that T_1__7 displayed mutations only in *SlDMR6-1* locus and to get a deep insight into possible nonspecific editing activity, we analysed the candidate off-target loci by using the resequencing data. At first we generated a list of 53 potential off-targets for the gRNA1, gRNA2 and gRNA3 used to target the *SlDMR6-1* locus (Data S1). All the 53 candidate off-target regions showed a number of mismatches higher than 2 bp with respect to the gRNAs. They fell in both non-coding (46) and coding (7) regions (Table 1).

**Table 1.** Analysis of *SlDMR6-1* off targets in plant T_1__7.

| gRNA | Number of off target in the genome | In coding | Non coding | SNP/Indel |
| --- | --- | --- | --- | --- |
| gRNA1 | 21 | 2 | 19 | 0 |
| gRNA2 | 22 | 4 | 18 | 0 |
| gRNA3 | 10 | 1 | 9 | 0 |
| Total | 53 | 7 | 46 | 0 |

An off-target analysis was conducted by aligning Illumina reads from both the WT and T_1__7 genomes to the tomato reference genome (Heinz 1706). This examination encompassed 53 potential off-target regions, ensuring a thorough evaluation to rule out the possibility of substantial deletions. Through a side-by-side comparison of DNA alignments between the WT and the mutant (*Sldmr6-1*), we ascertained that none of the candidate off-target regions exhibited any SNPs, indels, or significant deletions. Even if minor indels or SNPs were present in the surrounding areas, they did not signify off-target effects for two critical reasons: i) these variations were conserved between mutants and WT, while they displayed polymorphism compared to the Heinz 1706 genome; ii) variations were located beyond the 20-base pair window associated with the gRNA-like sequence. Our analyses unequivocally establish the specificity of Cas9-mediated *SlDMR6-1* gene editing, as evidenced by the absence of off-target effects.

Subsequently, leveraging resequencing data, we identified polymorphisms in both T_1__7 and WT, employing the Heinz tomato genome as reference. In T_1__7, we detected 42,196 SNPs, with 88.5% of them in a heterozygous state, while in WT, 40,998 SNPs were observed, of which 91.3% were in a heterozygous state. The average number of SNPs in both not edited plants (52.4 SNPs per Mb) and edited (53.9 SNPs per Mb) proved comparable, as did the average mutation rate (0.0054% for edited plants and 0.0052% for unedited plants), as documented in Table 2.

**Table 2.** SNPs statistics of WGS. WT and T_1__7 plants were compared at genomic level with reference genome (Heinz).

| Genotype | Plant type | SNPs | Homozygous | Heterozygous | SNP (%) | SNP per Mb |
| --- | --- | --- | --- | --- | --- | --- |
| T<sub>1</sub>_7 | edited | 42,196 | 4,86 | 37,336 | 0.0054 | 53.93 |
| WT | *in vitro* | 40,998 | 3,566 | 37,432 | 0.0052 | 52.40 |

### Effects of *SlDMR6-1* knock-out on drought resistance

To investigate the role of *SlDMR6-1* in drought stress resistance, six-week-old WT and T_2__6 and T_2__7 were subjected to drought stress conditions by withholding water during a further week. At the end of the 7 day–stress period WT plants exhibited severe wilting whereas T_2_ plants remained turgid whereas (Fig.3; Fig.S2).

**Fig. 3.**
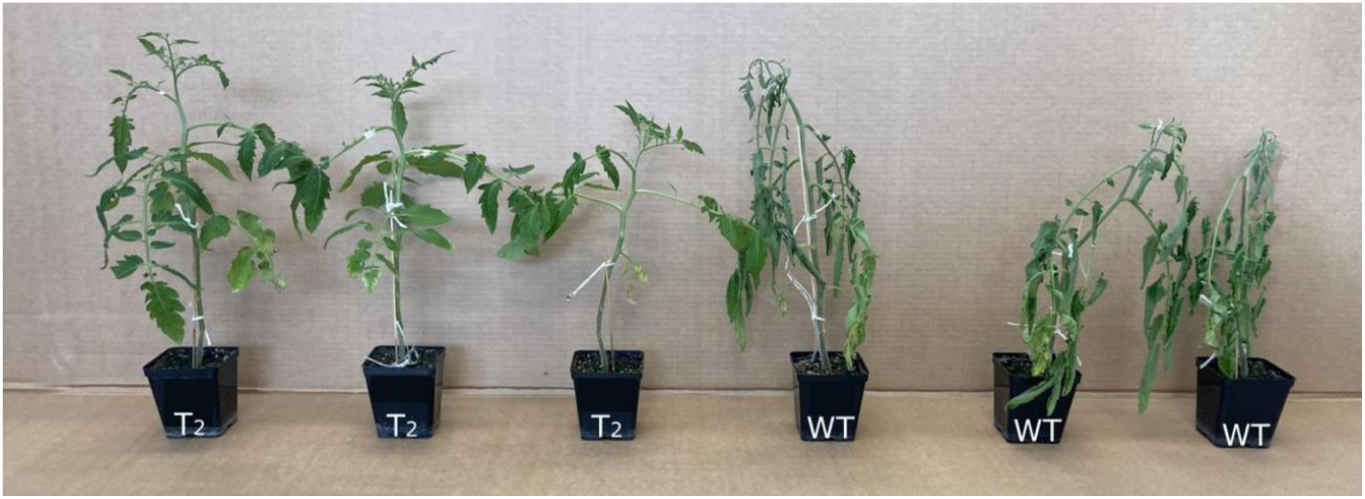
Drought stress analysis. T_2__7 *Sldmr6-1* plants and WT plants growing in a greenhouse after 7 days of withholding water

As shown in Fig.4, the Soil Relative Water Content (SRWC) of both the WT and the T_2__7 plants decreased during drought treatments, as expected. However, every day the rate of water loss in the T_2__7 plants was lower than that of WT, indicating that edited plants transpired less. Different lines of edited plants behaved similarly: although T_2__7 was slightly superior to T_2__6, no statistically significant differences between the two T_2_ lines were observed (Fig.S3).

**Fig. 4.**
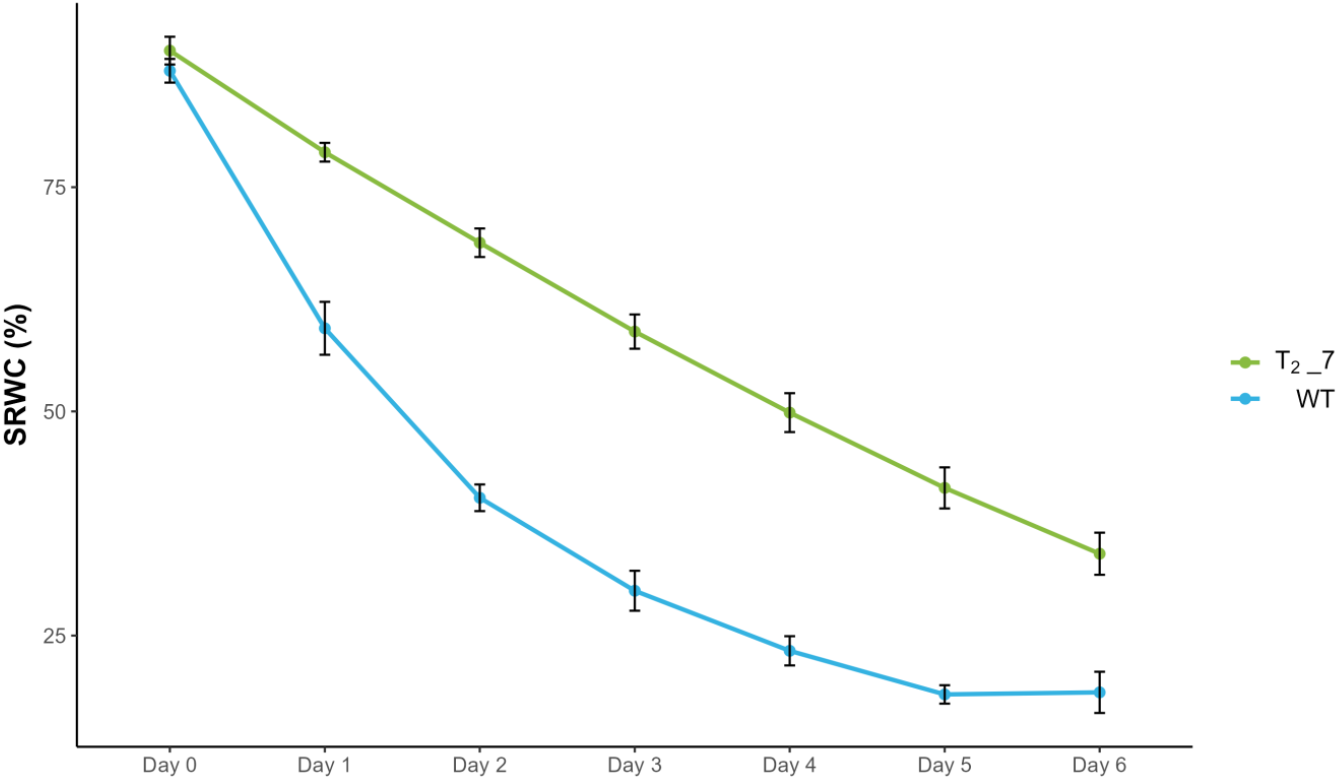
Soil relative water content (SRWC) of WT and *Sldmr6-1* lines (T_2__7) during the drought period. Each value represents the mean of six biological replicates ± SE.

Also, leaf area and dry weight (leaves, stems, roots) were evaluated highlighting no significant differences (Fig.S4).

During soil drying kinetics, assimilation rate (A), transpiration rate (E), stomatal conductance (Gs) and CO_2_ concentration in the sub-stomatal chamber (Ci) were measured in both WT and mutants, together with petiole water potential (Fig.5).

**Fig. 5.**
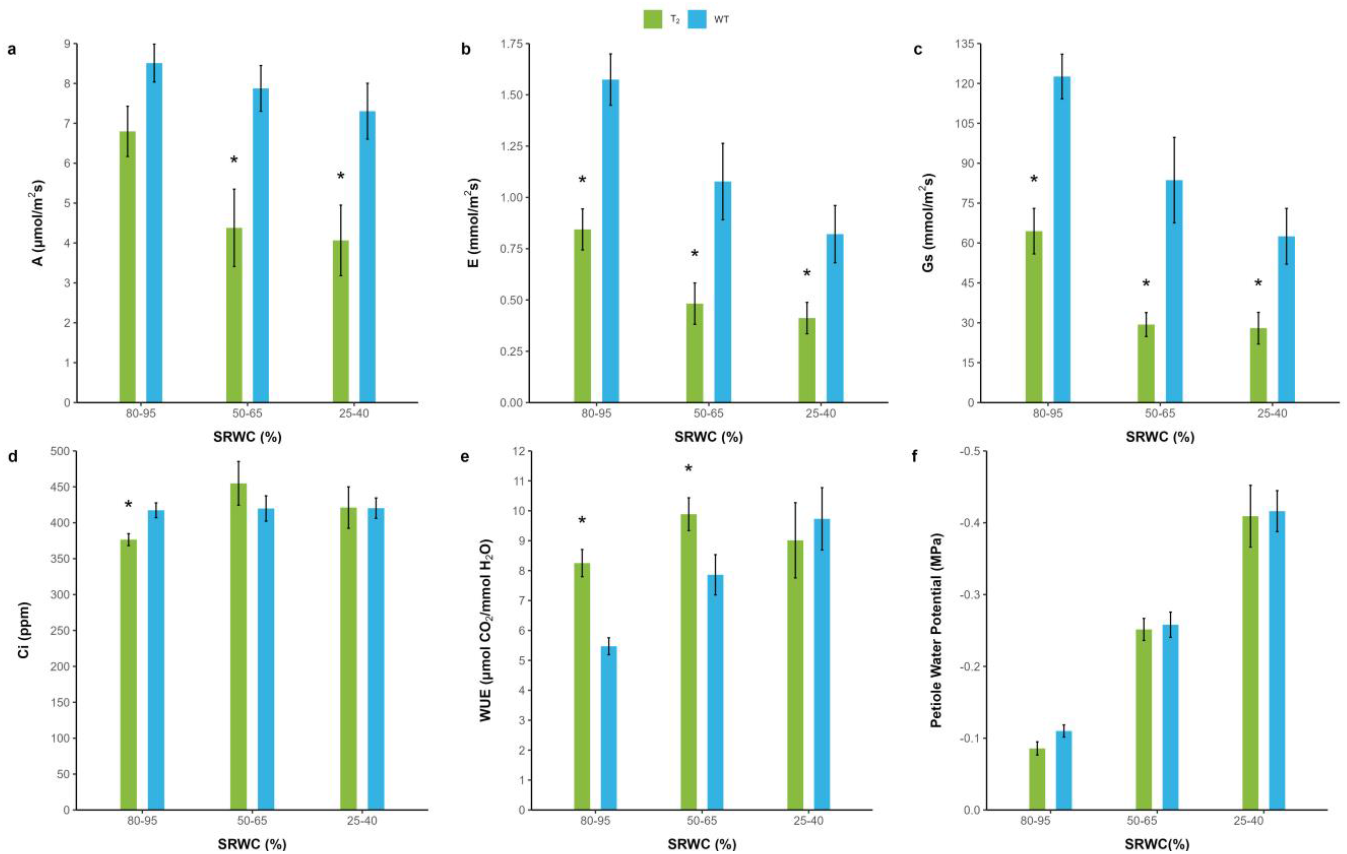
Leaf gas exchange of WT and *Sldmr6-1* lines (T_2__7) during the drought period, according to the decreasing trend of soil relative water content (SRWC). Plants under water stress were analysed to determine different eco-physiological traits: (a) assimilation rate (A), (b) transpiration rate (E), (c) stomatal conductance (Gs), (d) CO_2_ concentration in the sub-stomatal chamber (Ci), (e) Water Use Efficiency (WUE) and (f) petiole water potential. The data presented are the average values from six biological replicates with the standard error (SE) indicated. An asterisk denotes a statistically significant difference as determined by an ANOVA test (p ≤ 0.05).

Water Use Efficiency (WUE) was calculated as A/E. E and Gs were significantly reduced in T_2__7 plants with respect to the WT at any SRWC range. A was significantly reduced at SRWC range 50-65% (moderate stress) and 25-40% (severe stress), whereas Ci did not show significant differences. WUE significantly increased in T_2__7 at SRWC range 80-95% (no stress) and 50-65%. Ecophysiological traits of WT plants had an abrupt collapse concurrently with wilting at day 6; at this time point SRWC for WT plant was around 20% while was higher in T_2__7 plants (around 40%). No significant differences in ecophysiological traits were underlined by comparing T_2__6 and T_2__7 (Fig.S3). In *Sldmr6-1* plants, despite under conditions of reduced stomatal conductance, no metabolic damage that affect carboxylation activity was conceivable and this resulted in a Ci trend similar to what measured in WT controls, and a gain in WUE till to a moderate stress condition.

The increased drought resistance of *Sldmr6-1* lines prompted us to examine whether the expression of genes involved in ABA biosynthesis (*SlNCED1, SlNCED2, SlNCED3*) and catabolism (*SlCYP707*.*A1, SlCYP707*.*A2, SlCYP707*.*A3*) was altered in the edited lines under drought conditions. Moreover, we examined the transcript levels of key anti-oxidant related genes (*SlGST, SlPOD, SlSOD, SlAPX1, SlCAT1*) (Fig.6).

**Fig. 6.**
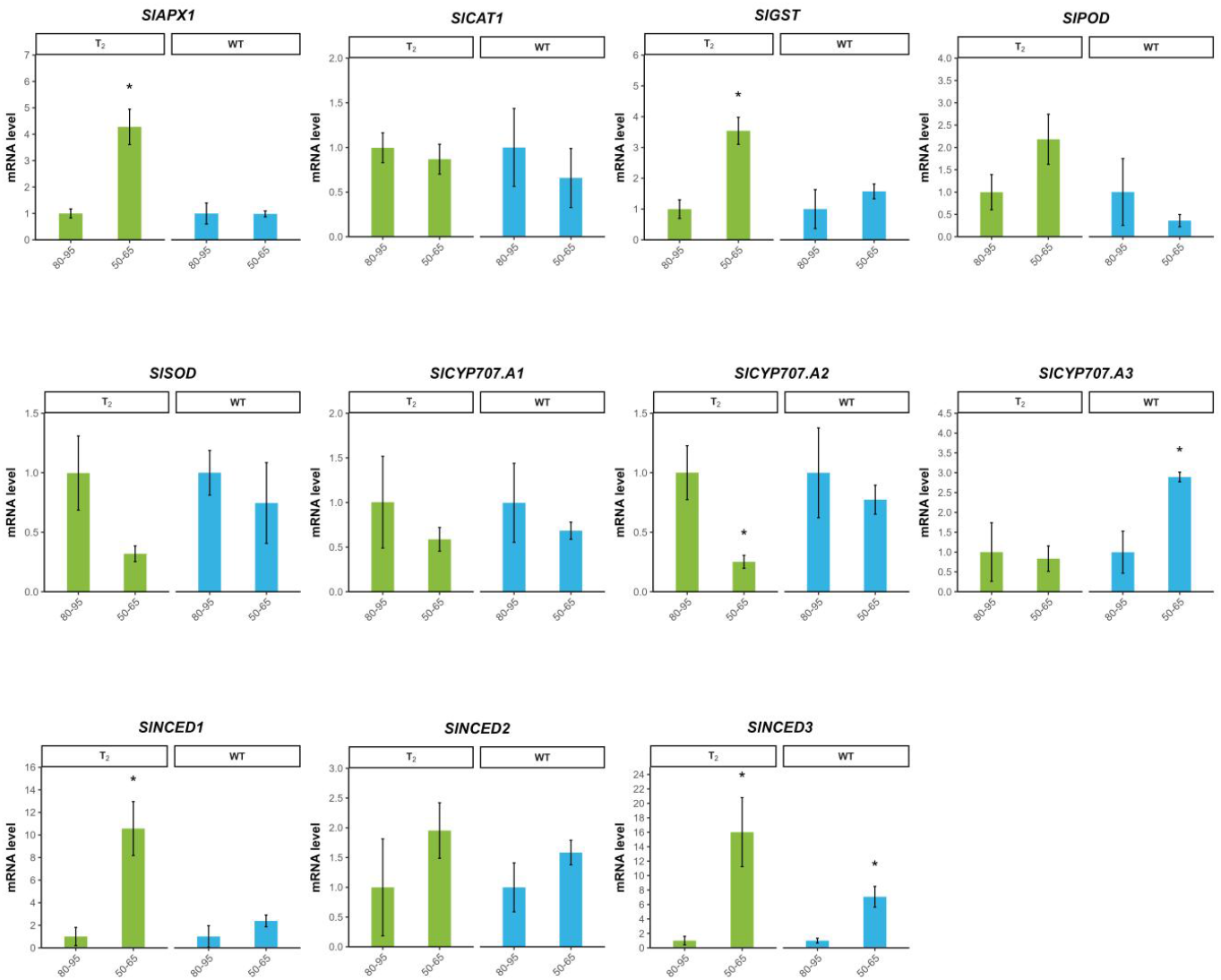
Transcriptional levels of anti-oxidant related genes (*SlAPX1, SlCAT1, SlGST, SlPOD, SlSOD*) and ABA-related genes (*SlCYP707*.*A1, SlCYP707*.*A2, SlCYP707*.*A3, SlNCED1, SlNCED2, SlNCED3*) during the drought assay. The values are expressed as relative mRNA abundance at SRWC range 50-65, and compared to SRWC range 80-95%. Tomato *actin* and *β*-*Tubulin* genes were used as reference genes. Data are means of three biological replicates ± SE. Data refer to T_2__7 line. An asterisk denotes a statistically significant difference as determined by an ANOVA test (p ≤ 0.05).

Among anti-oxidant related genes, a significant up-regulation in T_2_ plants was detected for *SlGST* and *SlAPX*. Among genes related to ABA biosynthesis, a strong up-regulation in T_2_ plants was demonstrated for *SlNCED1* and *SlNCED3* (around 10 and 16 fold higher respectively).

Among genes related to ABA catabolism, *SlCYP707*.*A2* was down-regulated in the T_2_ line, while *SlCYP707*.*A3* up-regulated in WT.

### Knock-out of *SlDMR6-1* improves tolerance against *P. infestans*

The impairment of S-genes leads to resistance or tolerance against several biotic stresses. *SlDMR6-1* knock-out in tomato is related to tolerance against a wide array of pathogens (De Toledo Thomazella et al, 2021). In this work we assessed tolerance against *P. infestans* in six selected T_1_ lines (T_1__6, T_1__7, T_1__11, T_1__12, T_1__13, T_1__14). A pathogenic assay was performed by using a detached leaf assay (Foolad et al, 2015). 72 hours after inoculation the edited T_1_ lines showed reduced susceptibility to *P. infestans* as highlighted by smaller necrotic and chlorotic foliar lesions than the control plants (Fig.7a). Genomic DNA was extracted from foliar disks cut around infection site and qPCR was used to quantify the fungal biomass (Fig.7b). Edited T_1_ lines showed a clear reduced fungal biomass, from 64% (T_1__6) to 95% (T_1__7, T_1__13) reduction compared to WT.

**Fig. 7.**
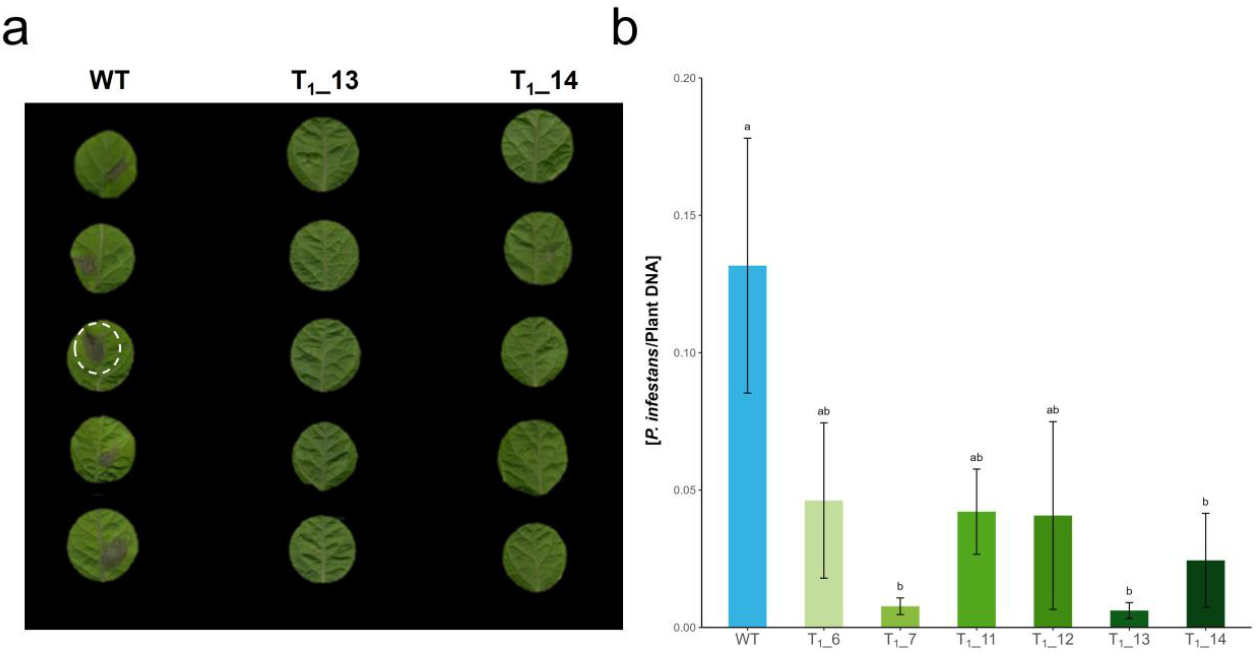
Pathogen assay on WT and *Sldmr6-1* lines (a) Detached leaves assay with *Phytophthora infestans* performed on two *dmr6-1* mutants and a WT plant as a control group at three days post-inoculation. The circle indicates the pathogen lesion (b) q-PCR pathogen DNA quantification after *Phytophthora infestans* infection. Data are the means of five biological replicates ± SE. Letters indicate significant differences based on Tuckey’s HSD Test.

## DISCUSSION

It has been previously reported that foliar application of salicylic acid (SA) to tomato plants under water deficit conditions can increase stomatal conductance, CO_2_ assimilation, and water use efficiency (WUE), mitigating the oxidative stress caused by ROS over-production (Aires et al, 2020). One of the key enzyme in SA metabolism is DOWNY MILDEW RESISTANT 6 (DMR6), which catalyzes the formation of 2,5-dihydroxybenzoic acid through the hydroxylation of SA at the C5 position of its phenyl ring (Zhang et al, 2017). Inactivation of *DMR6* results in increased SA acid levels (De Toledo Thomazella et al, 2021; Zeilmaker et al,2015).

Here we explored drought tolerance of *Sldmr6-1* tomato mutants obtained through CRISPR/Cas9. T_1_/T_2_ lines, characterized by the absence of any transgene and the disabling of *SlDMR6-1* locus in homozygosity were selected (Fig.1 and 2).

We examined the impact of the *Sldmr6-1* mutation on 7 days of water deprivation, and while the WT plants exhibited severe wilting, *Sldmr6-1* mutants showed turgid and green leaves (Fig.3). Under water deprivation, plants can adopt the strategy of modulating gas exchange by reducing the stomatal conductance and transpiration, resulting in lower assimilation of CO_2_ (Bucley et al, 2019). Ecophysiological traits measured during the period of water stress showed that the modification of *Sldmr6-1* prompted a water saving behaviour reducing Gs, and in turn E and A, and supporting an efficient photosynthetic metabolism, since no difference in Ci and an increase in WUE were observed (Fig.5).

The regulation of stomatal closure and maintenance of high soil Relative Water Content (SRWC) is an important strategy for water conservation under drought stress. In our study *Sldmr6-1* lines maintained higher soil SRWC than control plants during the whole imposed water stress of 1-week (Fig.4), presumably by reinforcing stomatal closure or preventing stomata opening.

Drought avoidance (referred to as dehydration avoidance in recent literature) occurs when plants increase their WUE by reducing transpiration and avoiding dehydration during periods of drought stress (Kooyers et al, 2015). The lower transpiration detected in our *Sldmr6-1* mutants suggests that their improved performance under deficit conditions was due to the drought avoidance mechanism, as previously observed in other tomato mutants (Shohat et al, 2021).

Water deficit causes the increase of reactive oxygen species (ROS) within plant cells, which provoke oxidative damage, especially in plants adopting drought avoidance strategy by reducing transpiration, which in turn increases the dangers associated with heating the leaves (Bleau et al, 2021). It has been previously demonstrated that the increased activity of enzymes such as peroxidase (POD), superoxide dismutase (SOD), catalase (CAT), glutathione S-transferase (GST) can contribute to the enhancement of drought resistance in tomato (Chen et al, 2021; Liang et al, 2022). Our data highlight that, following drought stress, *Sldmr6-1* mutants up-regulated the transcription of *SlGST* and *SlAPX* (Fig.6), two key anti-oxidant genes significantly upregulated when tomato plants were exposed to abiotic stress (Khan et al, 2015). The increased antioxidant activities in *Sldmr6-1* mutants might lead to a less severe oxidative damage under drought stress. The successful coupling between the drought avoidance strategy and an efficient ROS scavenging activity allowed stomatal control of photosynthesis, which did not cause metabolic imbalances causing negative feedbacks on photosynthetic activity. This is clearly evident from the overlapping of the Ci trends calculated on the basis of gas exchange data in both edited and WT plants and leading to the maintenance of high WUE values in the edited plants.

In response to water stress, a crosstalk between jasmonate acid (JA), SA, and abscisic acid (ABA) in tomato has been underlined (Muñoz-Espinoza et al, 2015) In salt-stressed tomato, SA modulated the expression of the genes involved in ABA accumulation and promoted the ABA transport to the shoot (Horváth et al, 2015). A cross-talk between ABA and SA signalling is thus important for the regulation of plant reproduction and growth under combined abiotic and biotic stresses (Horváth et al, 2015).

Drought avoidance is mainly regulated by ABA which induces stomatal closure by regulating the expression of many stress-responsive genes. The accumulation of ABA is regulated by a balance between its biosynthesis (catalysed by 9-cisepoxycarotenoid dioxygenase enzymes) and catabolism (catalysed by 8’-hydroxylases). For its synthesis, three *SlNCED* genes have been characterized in tomato, while for ABA catabolism, the *SlCYP707*.*A1, A2, A3*, and *A4* genes play a pivotal role (Liang et al, 2022). The contribution of *SlNCED1* or *SlCYP707*.*A2* in ABA accumulation has been demonstrated through over-expression and silencing approaches (Sun et al, 2012; Chatfield et al, 2000; Sun et a 2012b).

Our qPCR analyses demonstrated that the knock-out (KO) of *SlDMR6-1* prompted the up-regulation of *SlNCED1* and *SlNCED3* and downregulation of *SlCYP707*.*A2* upon stress application in contrast with WT plants. Contrary to what observed in *Sldmr6-1* KO lines, *SlCYP707*.*A3* was up-regulated in the WT (Fig.6). The results suggested that *Sldmr6-1* mutation increased endogenous ABA level by promoting ABA synthesis and suppressing ABA degradation, thereby positively affecting the stress resistance mechanisms of plants.

On the basis of our physiological data we can state that a higher SA content in *SlDMR6-1* KO lines might reduce ABA degradation, leading to drought avoidance through stomatal closure and reduced water loss in response to drought stress.

Although the KO mutation of *SlDMR6-1* has been demonstrated to confer a broad-spectrum disease-resistance phenotype in tomato (De Toledo Thomazella et al, 2021), the potential resistance to *Phytophthora infestans* (the causal agent of Late Blight) has never been tested. Late blight is a serious disease that may devastate an entire unprotected tomato crop within 7–10 days of infection. For the first time, our results showed an improved tolerance to Late Blight in *Sldmr6-1* tomato edited lines (Fig.7) in agreement with what was observed in potato (Kieu et al, 2021).

Even if the CRISPR/Cas9 approach induces, at target loci, random mutations that are functionally equivalent to spontaneously occurring ones, it is not always easy to predict the functional equivalence between natural and induced mutations. It has been suggested that most untargeted variations in edited lines are induced by somaclonal variation during *in vitro* culture, inheritance from the maternal plants and pre-existing variation across the germline (Sturme et al. 2022). Whole genome sequencing (WGS) can be a useful resource to assess the substantial equivalence of edited lines with their WT equivalent, because it provides comprehensive information about genomic variations, such as indels, SNPs and other structural differences. Several studies employed WGS analysis of WT and CRISPR/Cas9-edited plants to investigate the specificity of genome editing, and observed that off-target mutations occur at a much lower level than background mutations, due to pre-existing/inherent genetic or/and somaclonal variations (Liu et al, 2022; Sturme et al, 2022; Tang et al, 2018; Li et al, 2019; Wang et al, 2021). In agreement with previous observations, targeted deep sequencing of *Sldmr6-1* KO line at putative 53 off-target loci once again confirmed the lack of off-target effects (Table 1). The average number of SNP across not edited and edited (52.4 SNPs per Mb vs 53.9 SNPs per Mb) was similar, such as the average mutation rate (0.0052% for unedited plants, 0.0054% for edited plants) (Table 2).

Our results confirm the high specificity of CRISPR/Cas9 in tomato, which represents one of the “cleanest” tools to introduce targeted mutations. Our edited line does not carry any foreign DNA sequences but only carries an insertion, completely indistinguishable from spontaneously occurring mutations.

## CONCLUSIONS

We demonstrated for the first time that *SlDMR6-1* gene knock-out could contribute to the development of new varieties resistant to drought stress through a water saving strategy. The drought-avoidance mechanism observed in our *dmr6-1* mutants might be related to a successful coupling between the drought prevention strategy and an efficient ROS scavenging activity allowing stomatal control of photosynthesis. These results suggested that the drought avoidance in our *Sldmr6-1* lines could be correlated with the activation of antioxidant genes, leading to a more efficient ROS scavenging that could prevent the damage associated with leaf heating, likely danger when transpiration is reduced. On the other hand, the *SlDMR6-1* KO lines might reduce ABA degradation, leading to drought avoidance through stomatal closure and reduced water loss in response to drought stress. In addition, our results add *P. infestans* to the list of pathogens to which *SlDMR6-1* gene knock-out can confer resistance (*P. syringae pv. tomato, X. gardneri, X. perforans, P. neolycopersici, P*.*capsici*) (De Toledo Thomazella et al, 2021).

Our results might be extended to other crops, being *DMR6* orthologs interesting targets for improving multi-stress tolerance through genome editing tools.

When discussing the regulation of genome edited plants, it is important to understand if the frequency of unintended DNA mutations deriving from NHEJ-mediated repair, as it happens in CRISPR/Cas9 events, differs from the other conventional breeding methods. On the basis of our genomic analyses we can state that CRISPR/Cas9 represents a precise tool to introduce targeted mutations, since our edited line carries an insertion that makes it completely indistinguishable from spontaneous mutants.

All the data generated in this study makes gene editing of *DMR6-1* a highly reliable and valuable target for tomato breeding. Hence, we intend to pursue additional research to conduct a more thorough evaluation of the effects of *SlDMR6-1* inactivation on tomato production in field conditions and assess its potential as a strategy for tomato breeding.

## MATERIALS AND METHODS

### Target identification and vector construction

Three gRNAs (Data S1) targeting the first three exons (Fig.S1) of *SlDMR6-1* (ID Solyc03g080190) were designed using the online tool CRISPR-P 2.0 (hzau.edu.cn). The transformation vector pDGB3_alpha1was assembled through a Golden Braid (GB) cloning system (Sarrion-Perdigones et al, 2011, Sarrion-Perdigones et al, 2013; Vazquez-Vilar et al, 2017; Maioli et al, 2020) following GB software-directed procedures (https://gbcloning.upv.es/). Within the vector, the expression of h*Cas9* and *NptII* was driven by the CaMV 35S and *nos* promoters, respectively, while the gRNAs were placed in a polycistronic gRNA array under the control of the *At*U6-26 RNA PolIII promoter and sgRNA scaffold/terminator.

### Plant material and genetic transformation

Seeds of the cultivar ‘San Marzano’ were provided by Agrion (www.agrion.it) and were maintained in the Germplasm Bank of DISAFA (University of Torino, Italy). Fifty tomato seeds were sterilized in 2.5% sodium hypochlorite soaking for 20’ and then rinsed in sterile water three times. Sterile seeds were placed on sterile germination medium (1/2 MS + 15 g/l sucrose + 8 g/l plant agar) in plastic boxes, that were kept at 25°C in the dark for 72h before being transferred to a day/night cycle of 16/8 hours. After 10 days, plantlets presented fully grown cotyledons that were used for plant genetic transformation.

The final vector *pDGB3_alpha1_Tnos:NptII:Pnos_U6-26:tRNA:gRNA1-2-3_P35S:hCas9:Tnos* was introduced by heat shock into the *Agrobacterium tumefaciens* LBA4404 strain. Bacteria inoculum was prepared as follows. On the first day, *A. tumefaciens* was cultured in MGL (Data S1) added with streptomycin 50mg/l and kanamycin 50mg/l and incubated at 28°C over night (ON). On the second day, an aliquot of the culture was inoculated (1:50) in TY (Data S1) supplemented with 200µM acetosyringone and incubated at 28°C ON. The OD_600_ was evaluated and the bacterial solution was diluted to a final OD_600_ of 0.10-0.15 in TY medium supplemented with 200µM acetosyringone. Cotyledons of the seedling were cut in pieces of about 0.5 cm, which were dipped in bacterial culture for 10’, blotted dry on sterile paper and placed for 48h on a co-culture medium in the dark. Callogenesis, shoot induction, elongation and rooting were obtained as previously described (Qiu et al, 2007). After regeneration, fully developed plantlets (T_0_ plants, Data S1) were transplanted to soil and acclimated to *ex vitro* environment. By selfing T_0__2 plant (selected on the basis of editing outcome, Data S1), 14 plantlets were obtained (T_1_ generation) (Data S1). T_2_ plants were obtained from selfing of T_1__6 (homozygous, Cas positive) and T_1__7 plant (homozygous, Cas free).

### Molecular Screening

Genomic DNA was extracted from T_0_/T_1_ plants’ leaves using E.Z.N.A.® Plant DNA Kit (Omega Bio-Tek, Norcross, USA). The screening for h*Cas9* presence was performed using primers reported in Data S1 by PCR using KAPA HIFI Taq (Kapa Biosystems, Boston, USA) with the following program: 95°C/3’, 30 cycles of 98°C/15’’, 60°C/20’’, 72°C/1’ and 72°C/5’.

Editing efficiencies were evaluated by PCR amplification of gRNA-targeted regions according to Maioli et al. (Maioli et al, 2022) (Data S1). PCR products were sequenced by Sanger method and chromatograms were analysed using the TIDE online tool (Brinkman et al, 2018).

### Whole Genome Sequencing and analysis

A T1 plant (T1_ 7), carrying a homozygous mutation in two target regions and hCas9 segregation, together with a wild type (WT) plant were whole genome sequenced with an Illumina sequencer (Illumina Inc., San Diego, USA). One μg of DNA was used to prepare short insert (length 350 bp) genomic libraries (Novogene, Hong Kong), which were sequenced with paired-end chemistry (2×150 bp). Cleaning of the raw reads was conducted using Scythe (v0.991, https://github.com/vsbuffalo/scythe) and Sickle (v1.33, https://github.com/najoshi/sickle). SRA files (Project: PRJNA846963), containing raw data was submitted to NCBI.

A de novo genome assembly was carried out using the MegaHit assembler (v1.2.9, available at https://github.com/voutcn/megahit). This assembly process involved specific parameters, including k-min = 27, k-max = 141, k-step = 10, disconnect-ratio = 0, and cleaning-rounds = 1. Subsequently, a Blast analysis was conducted on the assembled scaffolds (T_1_ and WT) to identify potential insertions, utilizing the T-DNA sequence as the query.

### Target, off-target analysis, and SNP statistics

To analyze the identified target genomic variants and allele frequencies, we employed CRISPResso2 (accessible at http://crispresso2.pinellolab.org (Clement et al, 2019)). Fastq reads were extracted within a 100 bp window around each gRNA. For the identification of potential off-target regions in the tomato genome (SL4.0), we utilized the CasOT script (available at https://github.com/audy/mirror-casot.pl). All designed gRNAs were considered as baits in a single-gRNA mode, adhering to the default PAM type (NGG = A) and specific permissible mismatches in both the non-seed (2) and seed (2) regions. Coordinates of all potential target and off-target genomic regions were intersected with the vcf file using the bedtools intersect command (accessible at https://bedtools.readthedocs.io) to eliminate monomorphic regions among edited and WT plants. The results were then inspected through custom bash scripts.

For the edited plant samples, clean reads were aligned to the tomato reference genome (SL4.0, available at https://solgenomics.net) using the Burrows-Wheeler Aligner (BWA, v0.7.17, accessible at https://sourceforge.net/projects/bio-bwa/files). The ‘mem’ command with default parameters was employed for this purpose. Subsequently, BAM files were processed and used for SNP calling through Samtools (v1.9-166-g74718c2) mpileup, utilizing default settings with the exception of the minimum mapping quality (Q = 20) and filtering out multimapping events (-q > 1). This process resulted in the generation of a vcf (variant call format) file.

### Drought stress analysis

Six WT, 6 T_2__6 and 6 T_2__7 plants were grown in a climate chamber (temperature 25°C, RH 60%, 16 h light: 8 h dark photoperiod cycle, light intensity of 300 µmol m^-2^ s^-1^ PPFD) in pots containing perlite and soil-substrate (van Egmond universele potgrond) 1:5 v/v (Fig.S2). A quote of this soil was used to determine the maximum water holding capacity of the pots (Patono et al, 2022). Plants were grown in a well-watered state by watering to field capacity (above 75% of soil relative water content SRWC, daily at 8 am) for 6 weeks prior to the experimental imposed drought. Starting the drought, plants were allowed to slowly experience water stress by withholding irrigation. The measurement of petiole water potential allowed to identify two levels of water stress: a moderate water stress, when petiole water potential reached -0.3 MPa (day 1 in WT plants, day 4 in mutants), and a severe stress when petiole water potential had reached -0.5 MPa (day 3 in WR plants, day 6 in mutants). Plants at day zero (well-watered conditions) showed approximately – 0.1 MPa in both WT and mutants according to (Secchi et al, 2013). At the end of the drought stress, leaves were detached from plants and scanned. Pictures obtained were analysed with Image J software for leaf area evaluation. Leaves, stems and roots of single plants were collected, dried separately and weighted according to (Huang et al, 2019).

Steady state measurements of plant-to-atmosphere gas exchange were conducted on replicate plants from 10:0 am to 02:00 pm with a portable Infra Red Gas Analyzer - IRGA (GFS-3000, Walz, Germany) on single leaves, under 300 ± 5 µmol m^-2^ s^-1^ light, adjusted by the additional IRGA light source (Patono et al, 2023). Assimilation rate (A), transpiration rate (E), stomatal conductance (Gs) and CO_2_ concentration in the sub-stomatal chamber (Ci) were calculated following von Caemmerer and Farquhar’s equations (Von Caemmerer et al, 1981); water use efficiency (WUE) was calculated as A/E.

Measurements were taken in watered condition (Day 0) and daily following drought stress application (Day 1-7). One leaf per plant was sampled at each measurement, frozen in liquid nitrogen and stored at -80°C. Pots’ weight was also measured daily and soil relative water content (soil SRWC) calculated as percentage of moisture in the soil compared to the maximum water holding capacity.

RNA was extracted from leaf samples (three biological replicates for each genotype) and using Spectrum Plant Total RNA Kit (Sigma Aldrich, Saint Luis, USA) following instruction. qPCR analysis was performed according to Maioli et al. (2020). Chosen targets belong to two groups: anti-oxidant related genes (*SlGST, SlPOD, SlSOD, SlAPX, SlCAT*) and ABA-related genes (*SlNCED1, SlNCED2, SlNCED3, SlCYP707*.*A1, SlCYP707*.*A2, SlCYP707*.*A3*). Tomato *Actin* and *β*-*Tubulin* were used as housekeeping genes. Information about primer sequences and target can be found in Data S1. Transcript levels were quantified through the 2^-ΔΔCt^ method. Each value represented the mean of three biological replicates compared using Student’s t-test (p ≤ 0.05).

### Pathogen assay with *Phytophthora infestans*

The isolate of *Phytophthora infestans* (Westerdijk Fungal Biodiversity Institute strain CBS 120920) was maintained in Cornmeal medium (Data S1) at 18°C in dark. *P. infestans* was inoculated on Rye Agar (Data S1) one week before pathogenic assay and kept at 18°C in the dark. The plate was then flooded with chilled tap sterile water and kept for 2-3h at 4°C to induce zoospore release. Subsequently, the plate’s liquid was passed through two layers of cheesecloth, and the quantity of zoospores was determined using a hemocytometer. The concentration was diluted to 2.5 x 10^4^ spores/ml (Karki et al, 2021).

A detached leaf assay was set up using 5 leaves from the six selected T_1_ plants showed in Fig.1 and WT plants according to the procedure described by Foolad et al (2015). The leaves were washed with sterile water, gently dried using sterile paper, and then positioned in plastic trays containing water agar (20 g/l). About 250 zoospores (10µl) were placed on each leaf. Plastic trays were then covered with lids and incubated at 20°C in the dark in a growth chamber. The trays were examined every day. Picture and samples were collected three days post inoculation.

To quantify the pathogen infection rate, the ratio between fungal and plant DNA was evaluated according to Pavese et al. (2021). Disk samples around infection site were taken and DNA extraction was performed using an E.Z.N.A.® Stool DNA Kit (Omega Bio-Tek, Norcross, USA). For the quantification of DNAs standard curves were prepared using primers designed as follows: *SlActin* for tomato DNA, *PiO8* (Llorente et al, 2010) for *P. infestans* DNA. Extracted DNAs were analyzed through real-time qPCR both with pathogen gene (*PiO8*) and tomato’s one (*SlActin)*. qPCR reaction was carried out as described in the previous paragraph and information about primer sequences can be found in Data S1. Fungal and plant DNA was quantified using standard curves and the ratio fungus DNA/plant DNA calculated. One-way analysis of variance test (ANOVA)was performed through IBM SPSS statistical software. Each value represented the mean of 5 biological replicates compared using Tukey’s HSD Test (p ≤ 0.05).

## ACCESSION NUMBERS

| Gene | Species | Accession Number |
| --- | --- | --- |
| *DMR6-1* | *Solanum lycopersicum* | Solyc03g080190 |
| *GST* | *Solanum lycopersicum* | XM_004246333 |
| *APX1* | *Solanum lycopersicum* | Solyc06g005160.2.1 |
| *SOD* | *Solanum lycopersicum* | Solyc06g048410.2.1 |
| *POD* | *Solanum lycopersicum* | Solyc01g006300.2 |
| *CAT1* | *Solanum lycopersicum* | Solyc12g094620.1.1 |
| *CYP707.A1* | *Solanum lycopersicum* | EU183406 |
| *CYP707.A2* | *Solanum lycopersicum* | AK326035 |
| *CYP707.A3* | *Solanum lycopersicum* | AK321536 |
| *NCED1* | *Solanum lycopersicum* | Z97215 |
| *NCED2* | *Solanum lycopersicum* | EU912387 |
| *NCED3* | *Solanum lycopersicum* | GQ222384.1 |
| *ACTIN* | *Solanum lycopersicum* | NM_001308447 |
| *TUBULIN* | *Solanum lycopersicum* | Solyc04g081490.2.1 |
| *O8-3* | *Phytophthora infestans* | GQ371195.1 |

## DATA AVAILABILITY STATEMENT

Sequencing data used in this study are openly available in the NCBI database (PRJNA846963).

## ONLINE RESOURCES INFORMATION

| File name | Description |
| --- | --- |
| Supplementary Figures | **Fig.S1.** Position of gRNAs' target sites on *SIDMR6-1*. |
|  | **Fig.S2.** T<sub>2</sub>_6 *Sldmr6-1* plants and WT plants under drought stress |
|  | **Fig.S3.** Comparative a) ecophysiological and b) SRWC measurement of T<sub>2</sub>_6 and T<sub>2</sub>_7 |
|  | **Fig.S4.** Comparison of a) leaf area and b) dry weight between WT and T<sub>2</sub>_7 line. |

