## Supplementary_figures for "New insights on the role of *SlDMR6-1* in drought avoidance in tomato"

**Supplementary Files**

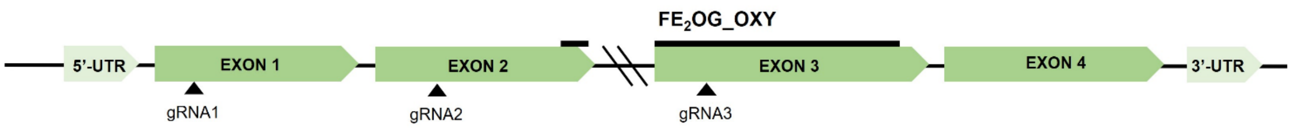


**Fig.S1** Position of gRNAs' target sites on *SlDMR6-1*

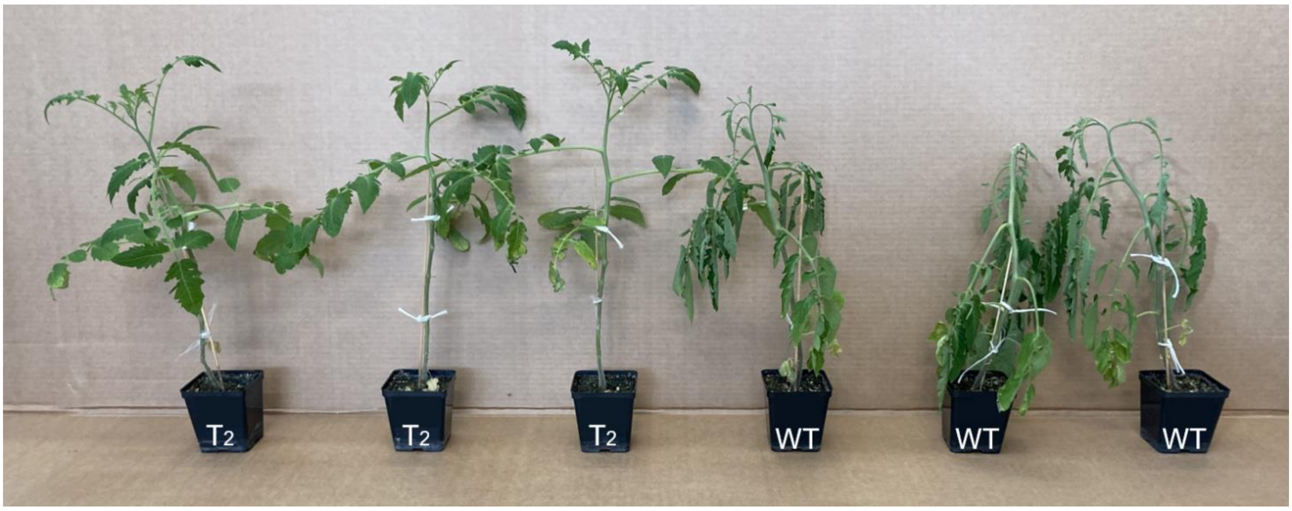


**Fig.S2** $T_2$_6 *Sldmr6-1* plants and WT plants under drought stress

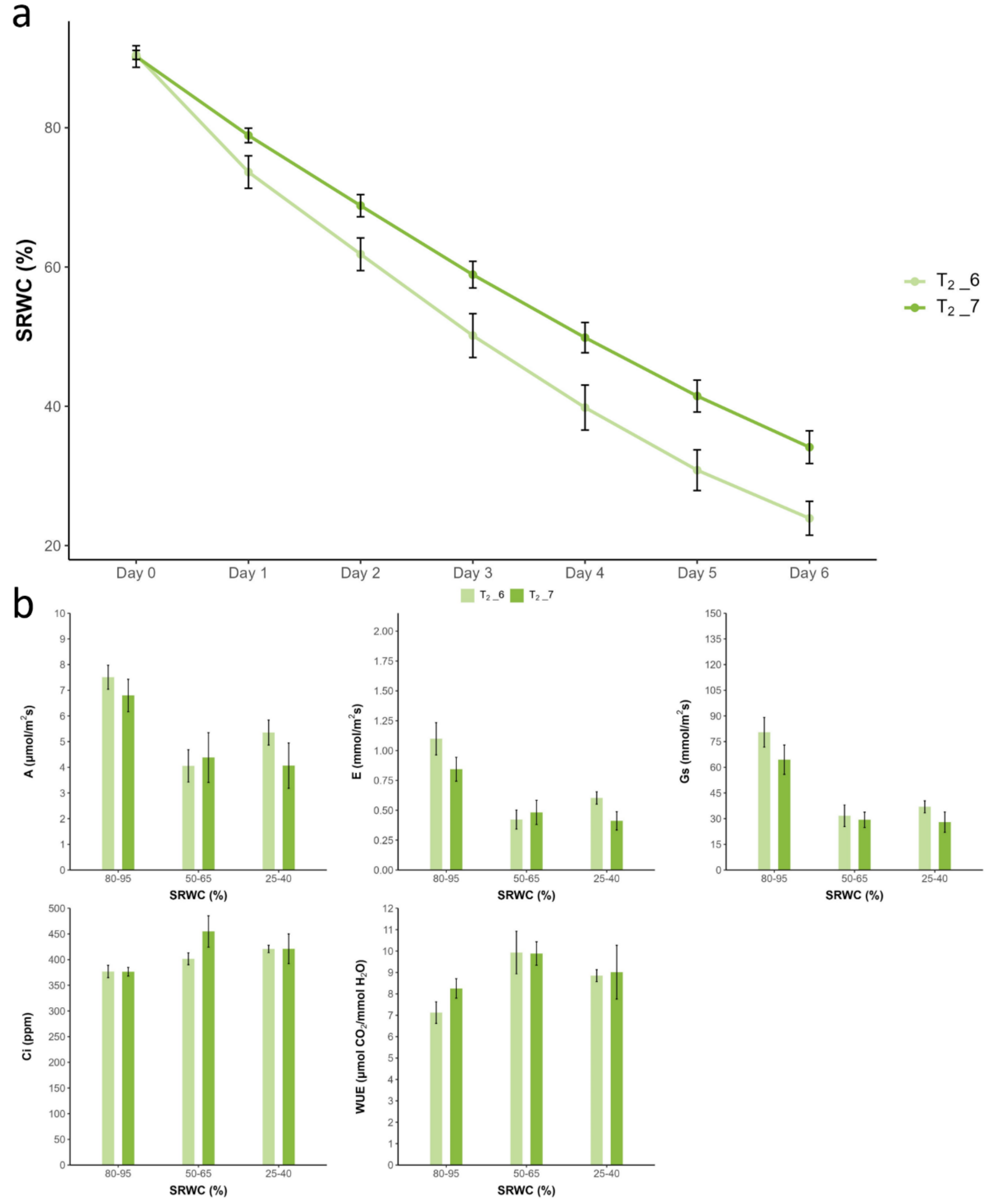


**Fig.S3** Comparative a) SRWC and b) ecophysiological measurement of $T_2$_6 and $T_2$_7

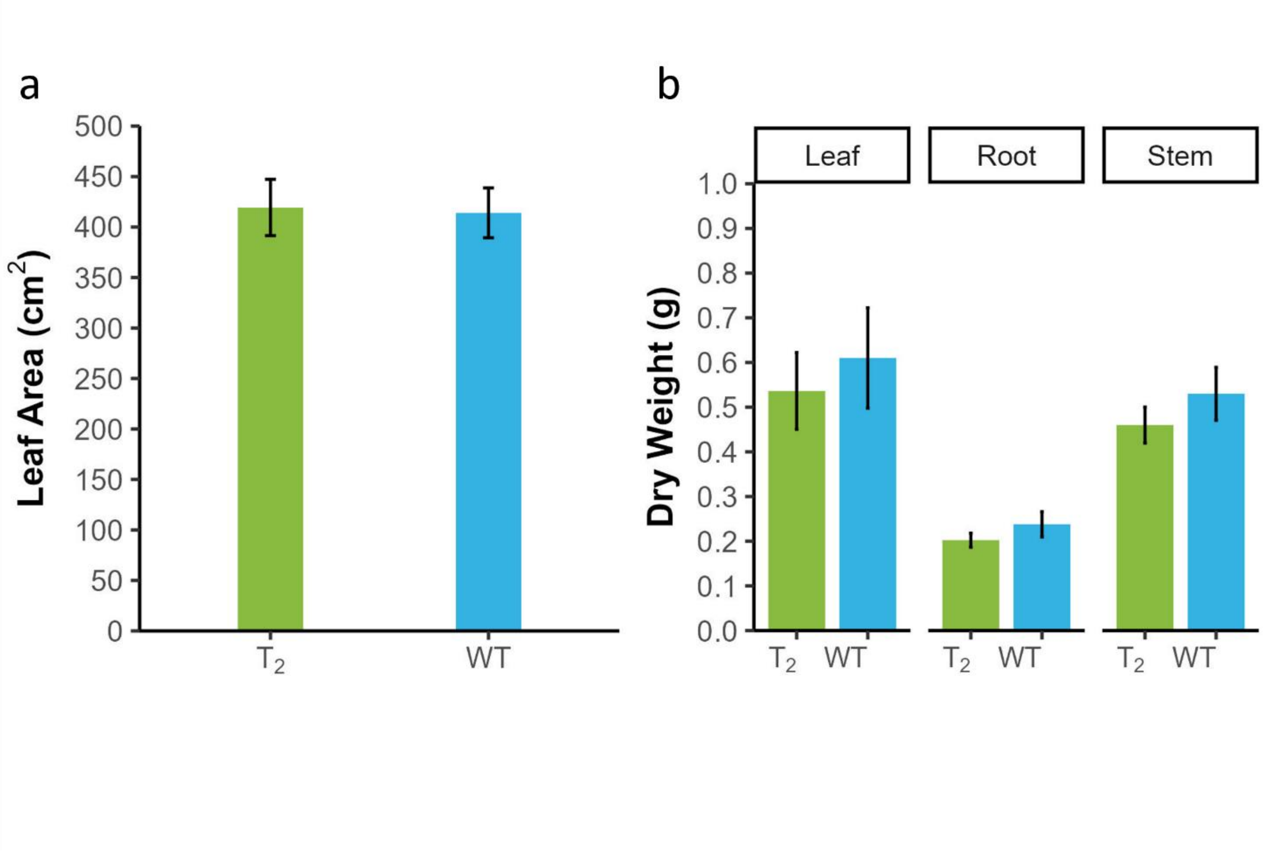


**Fig.S4** Comparison of a) leaf area and b) dry weight between WT and $T_2$_7 line
